# Follistatin-like protein 1 interacts with programmed cell death 4 to promote vascular mimicry in hepatocellular carcinoma by activating the PI3K/AKT/mTOR signaling pathway

**DOI:** 10.1101/2025.04.30.651401

**Authors:** Zhaowei Yang, Jiajing Zhu, Huan Chen, Jin Liu, Wenjia Tan, Hang Li

**Affiliations:** Department of Laboratory Medicine, China-Japan Union Hospital of Jilin University, Changchun, Jilin Province 130033, P.R.China; Department of Radiology, China-Japan Union Hospital of Jilin University, Changchun 130033, Jilin Province, P. R. China; Woman and Children’s Health Hospital of Jilin Province,130061, P.R.China; Department of Ultrasound, China-Japan Union Hospital of Jilin University, Changchun, Jilin Province 130033, P.R.China; Department of Hepatobiliary Pancreatic Surgery,China-Japan Union Hospital of Jilin University, Changchun 130033, Jilin Province, P. R. China

## Abstract

Hepatocellular carcinoma (HCC), the most common type of liver cancer, is highly prone to recurrence and metastasis after curative treatment. However, the molecular mechanisms governing these post-management setbacks remain largely unknown. Previous studies found that the high expression of follistatin-like protein 1 (FSTL1) in tumors is implicated in recurrence, metastasis, and poor patient prognosis. In this study, we aimed to investigate the role of FSTL1 in HCC.

We found that FSTL1 overexpression was positively correlated with vascular mimicry and poor prognosis. In vitro experiments confirmed that the interaction between FSTL1 and programmed cell death 4 (PDCD4) activates epithelial-to-mesenchymal transition and promotes the proliferation, invasion, and migration of liver cancer cells. The protein-protein interaction between FSTL1 and PDCD4 significantly promotes VM as well as the expression of the pro-angiogenic vascular endothelial growth factor by activating AKT/mTOR signaling. Our data suggest that FSTL1 may be a potential target for the treatment of HCC.

Implications: Our findings indicate that the protein FSTL1 may be a novel therapeutic target for managing patients with HCC.

## Introduction

Hepatocellular carcinoma (HCC) is a common type of malignancy that afflicts individuals worldwide; moreover, it is the third most common cause of cancer-related death (1). Early metastasis is a hallmark of the frequent recurrence and high mortality among patients with HCC; however, the molecular mechanisms responsible for this tumor’s aggressiveness remain unexplored. Epithelial-to-mesenchymal transition (EMT) is an important precursor for tumor metastasis (2). Follistatin-like protein 1 (FSTL1), also known as TSC-36 or FRP (3, 4), was originally isolated from the mouse osteoblastic cell line MC3T3E1 and is upregulated via stimulation by transforming growth factor-β1. Previous studies have shown that FSTL1 is involved in organ development (5, 6), inflammation, stem cell regeneration (7), and tumor development (8, 9) by mediating cell survival, proliferation, migration, and metastasis. In contrast, other studies have found FSTL1 to exert anti-tumor functions in nasopharyngeal (3), kidney, ovarian (10), and endometrial cancers (3). As such, the function of FSTL1 in tumor progression appears to be complex and not fully understood. Unlike angiogenesis in the classical sense, vascular mimicry (VM) supplies blood to tumor cells independent of endothelial cells (11). VM is associated with high tumor grade as well as tumor progression, invasion, and metastasis, and portends a poor patient prognosis (12, 13). In this study, we found that FSTL1 is significantly overexpressed in HCC tissues, and sought to investigate whether this attribute is related to this malignancy’s aggressive phenotype. We also aimed to clarify the signaling pathways associated with FSTL1 activity in HCC.

## Materials and Methods

### Clinical samples

HCC tissue specimens (n = 160) were obtained from the China-Japan Union Hospital of Jilin University (Jilin, China). The specimens were utilized for research purposes following ethical approval (23/09/2020), with data analysis completed by 06/12/2024. The patients had not undergone any treatment at the time of biopsy, and all had received a diagnosis of HCC upon pathological analysis of the tissue. The pathological diagnosis, pathological stage, and clinical data of all patients included in this study were collected from inpatient medical records, and survival data were collected via telephone follow-up. Ten control tissue specimens were acquired from healthy individuals who had visited the Physical Examination Center of China-Japan Union Hospital of Jilin University. Written informed consent was obtained from all participants in both the experimental and control groups. The TNM classification was determined according to the American Joint Committee on Cancer 7th edition criteria. The research involving these tissue samples was approved by the Ethics Committee of the China-Japan Union Hospital of Jilin University.

### Immunohistochemical staining

The tumor tissues were fixed in formalin, embedded in paraffin, and sectioned. Next, antigen retrieval was performed following which endogenous peroxidase was quenched. Slides were blocked with 10% donkey serum for 1 h at room temperature and then incubated with primary antibody overnight at 4°C. The slides were then washed with phosphate-buffered saline and exposed to the secondary antibody for 2 h at room temperature. The slides were washed again with phosphate-buffered saline, and colorimetric staining was visualized by incubating with DAB for no longer than 5 minutes (the reaction was stopped when the color intensity was moderate). The samples were then stained with hematoxylin for 3 minutes, dehydrated, and mounted. The percentages of positively-stained cells were then determined.

### CD31/periodic acid-Schiff (PAS) double staining and VM analysis

After immunohistochemical staining for CD31, the sections were washed with distilled water and incubated with periodic acid solution for 7 min. Next, the slides were exposed to Schiff’s solution for 20 min, after which the sections were washed with distilled water for 3 min and stained with hematoxylin. Gradient dehydration was then applied prior to mounting with neutral gum. Endothelium-dependent vessels were evaluated by counting CD31-positive vessels. The average numbers of PAS-stained and endothelium-dependent vessels at the peripheral zones of tumor necrosis were calculated using 10 randomly selected fields. In terms of evaluating VM, the presence of red blood cells in tumor cell-surrounded lumens that lacked vascular endothelial cells was considered VM-positive, while the absence thereof was considered VM-negative. The red blood cells were separated from tumor cells by continuous or discontinuous PAS-positive membranes with no obvious necrosis or inflammatory cell infiltration.

### Western blotting

Expression levels of FSTL1, programmed cell death 4 (PDCD4), VIM, E-cadherin, Snail, vascular endothelial growth factor (VEGF), hypoxia-inducible factor 1⍺ (HIF-1⍺), phosphatidyl inositol-3 kinase (PI3K), phospho-PI3K, Akt, phospho-Akt, mTOR, and phospho-mTOR were assessed using western blotting. Cell lysate proteins were separated using SDS-PAGE and transferred onto polyvinylidene difluoride membranes for probing. The details of the primary antibodies used are as follows: rabbit anti-FSTL1 (1:500, Proteintech, Rosemont, IL, USA), mouse anti-VIM (1:20,000, Proteintech), rabbit anti-PDCD4 (1:1,000, Abcam, Waltham, MA, USA), rabbit anti-E-cadherin (1:1,000, Abcam), rabbit anti-Twist1 (1:1,000, Abcam), rabbit anti-Snail (1:1,000, Abcam), rabbit anti-phospho-PI3K (1:1,000, Abcam), rabbit anti-PI3K (1:1,000, Abcam), rabbit anti-Akt (1:500, Abcam), rabbit anti-phospho-Akt (1:1,000, Abcam), rabbit anti-mTOR (1:2000, Abcam), and rabbit anti-phospho-mTOR (1:1,000, Abcam). The housekeeping protein GAPDH (1:2,000, Proteintech) was used to normalize protein content. After incubating the membranes with a fluorescent dye-labeled secondary antibody, the membranes were visualized and the densities of the bands were quantified using ImageJ (NIH, Bethesda, MD, USA) to calculate the relative protein expression levels.

### Quantitative real-time (qRT) PCR

Total RNA was isolated using the TRIzol reagent (Invitrogen, Waltham, MA, USA) according to the manufacturer’s instructions. First-strand cDNA was synthesized using a PrimeScript RT Reagent Kit (TaKaRa Bio Inc., Changchun, China). qRT-PCR was performed using the SYBR Premix Ex Taq Kit (TaKaRa Bio Inc.) on an Applied Biosystems 7500 Real Time PCR system (Applied Biosystems, White Plains, NY, USA). GAPDH was used to correct for gene copy size. The sample reaction was performed in a short time in a low temperature environment; data are reported as the fold changes. The primers used to probe each target gene are shown in Table 1.

**Table 1.**
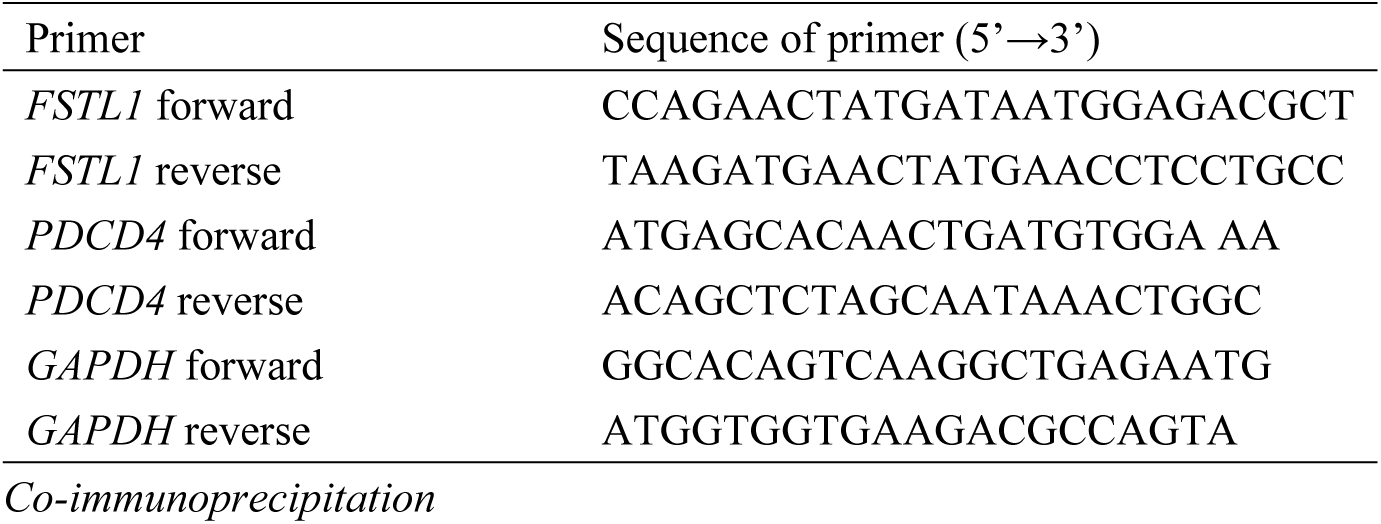
Primers used for quantitative real-time PCR.

### Co-immunoprecipitation

HCC cells transfected with *FSTL1*-Flag were lysed in RIPA buffer and then loaded onto Flag antibody-bound protein A/G agarose beads (sc-2021, Santa Cruz Biotechnology, Dallas, TX, USA). After incubation, the beads were washed 5 times with RIPA buffer.

### Cell lines, culturing, and transfection

The HCC cell lines HepG2, MHCC-97H, and Bel7402 were purchased from Keygene BioTECH (Shanghai, China). HepG2 and MHCC-97H cells were cultured in DMEM (Invitrogen) containing 10% fetal bovine serum (Gibco, Waltham, MA, USA) and 1% penicillin-streptomycin (Invitrogen). Bel7402 cells were cultured in RPMI-1640 (Invitrogen) containing 10% fetal bovine serum and 1% penicillin-streptomycin (Invitrogen). All three HCC cells were cultured in a 37 °C incubator with 5% CO2.

FSTL1 and PDCD4 overexpression plasmids as well as FSTL1 small hairpin (sh) RNA plasmids were constructed by Sangong Biotechnology (Shanghai, China); these were labeled with green fluorescent protein (GFP). The knockdown sites’ sense sequences were as follows:

Sh/FSTL1-1 GCTAGAGCCGCAAATTGCAGT, Sh/FSTL1-2 GTGTATAGTGCCAATCAGAAA. Two control constructs, OvCtrl and Sh/SiCtrl, were transfected using the empty vector pEGFP-N1 and pSGU6/GFP/Neo, respectively. Lipo2000 (Invitrogen) was used for transfection, and stably transfected cell lines were selected using kanamycin and by employing the limiting dilution method.

### Cell Counting Kit-8 (CCK-8) assay

Cells were extracted during their growth phase and their concentration was adjusted to 2×104/mL; 100 µL of this solution was plated into each well of a 96-well plate. At 0, 24, 48, and 72 h, 10 µL of CCK-8 (Beyotime Inst Biotech, China) was added to each well; the plate was gently shaken and incubated at 37°C for 2 h. The resulting solutions’ absorbances were measured using a microplate reader.

### Colony formation assay

Cells (100 per well) were seeded into 6-well plates and allowed to grow for 2 weeks. The formed colonies were fixed and stained with 0.1% crystal violet for 30 min; they were then examined under a microscope and counted for statistical analysis.

### Wound healing and transwell migration assays

HepG2 and MHCC-97H cells that were growing in the logarithmic phase were plated in a 6-well plate. After 12 h, the adhered cell monolayer in each well was scratched in a straight line. The scratched areas were then photographed at 24, 48, and 72 h and their widths were measured digitally. Matrigel-coated transwell chambers (catalog #3422, Corning Inc., Corning, NY, USA) were used to assess cell invasion in vitro. One hundred microliters of HCC cells suspended in serum-free media at appropriate concentrations were introduced into the upper chamber of each transwell, while 600 µL of complete medium containing 10% fetal bovine serum was added to the lower chamber. The upper chambers were removed after 12 or 24 h of incubation at 37°C in a 5% CO2 environment, and cells that had traversed the Matrigel to the bottom chamber were fixed, stained, and photographed under a microscope.

### Three-dimensional culture assay

Matrigel (Corning, Tewksbury, MA, USA) was liquified at 4°C, and 100 µL of the product was applied to each well of a 48-well plate using a pre-chilled pipette tip; the Matrigel was then allowed to gel at 37°C for 30 min. HCC cells were suspended at the appropriate concentrations and seeded onto the Matrigel, incubated for 6 hours at 37°C in a 5% CO2 environment, and photographed. The lengths of the formed vessels were measured using the ImageJ software (NIH).

### Xenograft model

All the animals used in our experiments were treated in accordance with current ethical standards and international conventions. MHCC-97H cells transfected with OvFSTL1, ShFSTL1, and control plasmids were injected subcutaneously into 5-week-old BALB/c nu/nu nude mice (Beijing Huafukang Biotechnology Co. Ltd., Beijing, China) while anaesthetized with 1% sodium pentobarbital. The long (a) and short (b) diameters of developing tumors as well as the animals’ weights were measured every 3 days, and the relative tumor volumes were calculated using the formula 0.5 × a × b × b. Mice were euthanized when the long diameter (a) reached 20 mm per our institutional guidelines on the ethical handling of animals.

### Statistical analysis

The data were analyzed using SPSS 19.0 (IBM Corp., Armonk, NY, USA) and are reported as means ± standard deviations. Student’s t-test was used to compare differences between 2 groups, while the Kaplan-Meier method was used for survival analysis. A P-value <0.05 was considered statistically significant.

### Ethics approval and consent to participate

Human study has been approved by Ethics Committee of China-Japan Friendship Hospital of Jilin University(2020-NSFC-074).The experiments confirm that informed consent has been obtained from all subjects and/or their legal guardians.The experiments conformed to the Guide for the Care and Use of Laboratory Animals.Animal study has been approved by the Institutional Animal Care and Use Committee of Jilin University(KT202402255). All methods are reported in accordance with ARRIVE guidelines.

## Results

### FSTL1 was overexpressed in human HCC samples and associated with VM and other clinicopathological features

To investigate the role of FSTL1 in HCC, we first evaluated FSTL1 expression in HCC and normal liver tissue samples using qRT-PCR and western blotting; we found that this protein was markedly upregulated in HCC (P<0.05) (Fig. 1A). We next evaluated FSTL1 expression in HCC using immunohistochemistry while simultaneously examining VM using CD31-PAS double-staining. VM channels were observed in 105 of 160 HCC samples, and were lined by tumor cells but lacked inflammatory cell infiltration or evidence of necrosis (Fig. 1). We categorized the 160 samples into 2 groups: FSTL1-positive (n=105) and FSTL1-negative (n=55); FSTL1 staining was observed in the cytoplasm (Table 2). VM was observed in both the FSTL1-positive and FSTL1-negative tissues (Fig. 1B); however, FSTL1 expression was associated with a higher probability of VM (Table 3). We also investigated the correlation between FSTL1 and pathological features, and found that FSTL1 was significantly correlated with histological differentiation, cancer stage, tumor size, and metastasis. Moreover, FSTL1 levels were higher in samples from patients aged ≥55 years than in those from patients <55 years, although the difference was not significant (P>0.05). FSTL1 expression was significantly higher in samples from patients with TNM stages III/IV than in those with stages I/II. However, FSTL1 expression was not associated with sex (P>0.05) (Table 2).We performed co-immunoprecipitation assays to determine whether FSTL1 interacts with PDCD4. PDCD4 in HCC cell lysates was precipitated by Flag-FSTL1-tagged beads but not by control beads following treatment with the mitochondrial uncoupling agent carbonyl cyanide m-chlorophenylhydrazone, demonstrating the binding of endogenous FSTL1 to PDCD4 in HCC cells(Fig.1D.

**Fig. 1.**
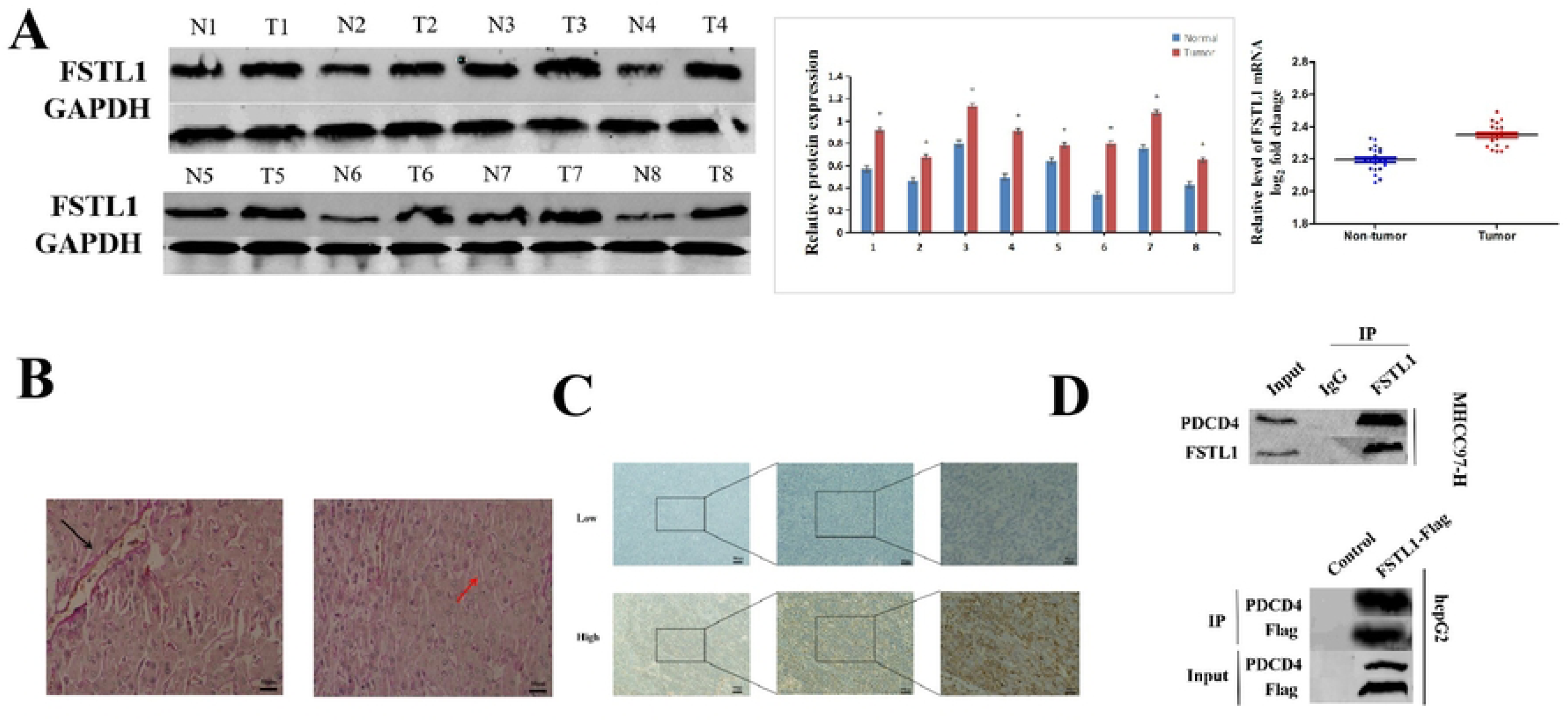
High suppressor of Follistatin-like protein 1 (FSTL1) expression predicts poor prognosis in hepatocellular carcinoma (HCC).**A** Western blotting and RT-PCR analysis of FSTL1 protein expression in HCC (T) and non-tumor liver tissues (N). FSTL1 protein expression levels were normalized according to GAPDH expression levels (n = 8 per group).**B** Evidence of VM (red arrow) and angiogenesis (black arrow) in HCC samples (400x, scale bar=50μm).**C.** Immunohistochemical analysis of FSTL1 protein expression in human HCC tissues.**D** .HCC cells were transfected with FSTL1-Flag and subjected to immunoprecipitation using anti-Flag mAb. Co-immunoprecipitated PDCD4 was detected using anti-PDCD4 antibody (up panel).Endogenous FSTL1 in HCC cells was immunoprecipitated using anti-FSTL1 antibody with IgG as nonspecifc control (down panel). Co-immunoprecipitated PDCD4 was detected using anti-PDCD4 antibody.

**Table 2.**
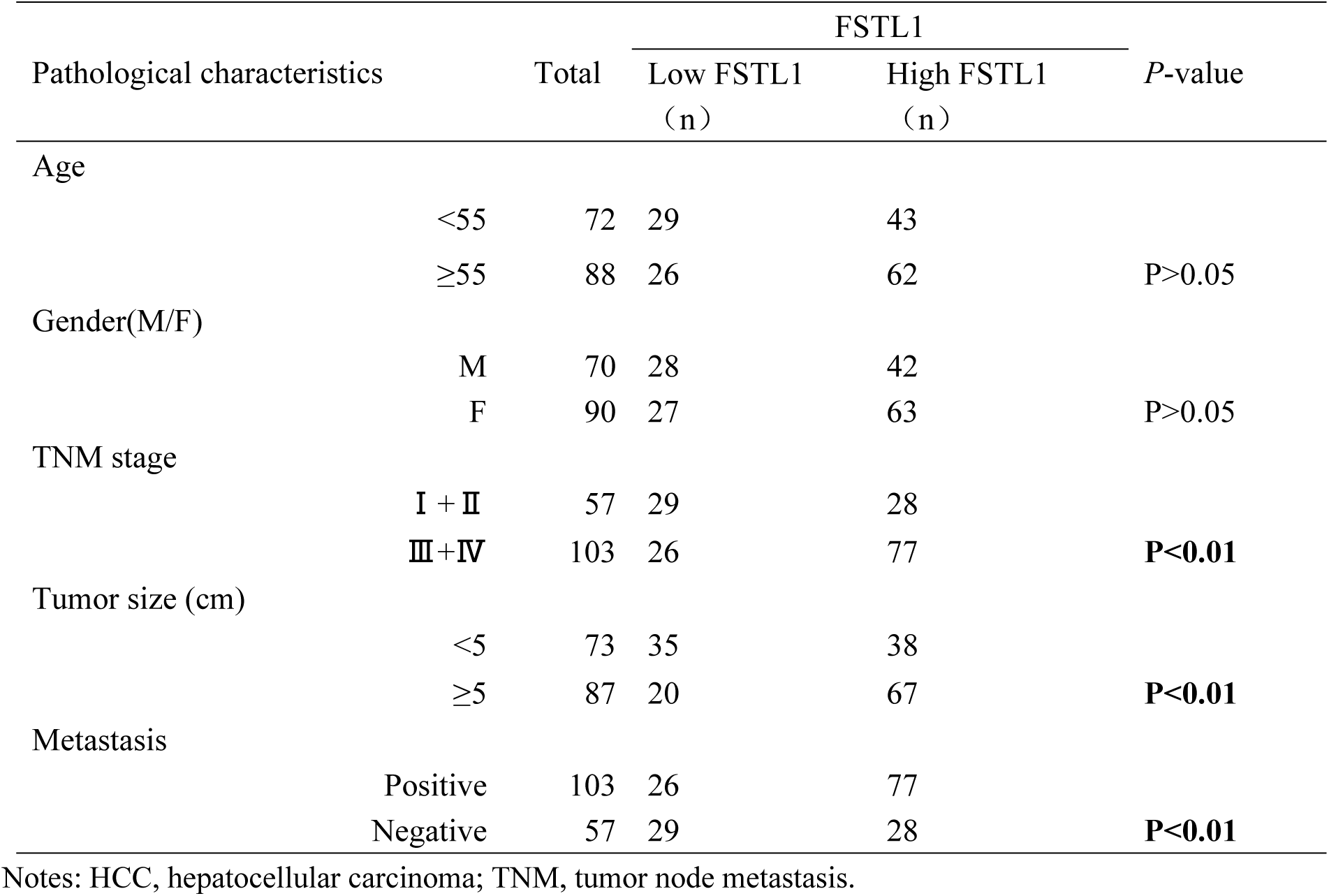
Characteristics of patients from whom hepatocellular carcinoma samples were extracted.

**Table 3.**
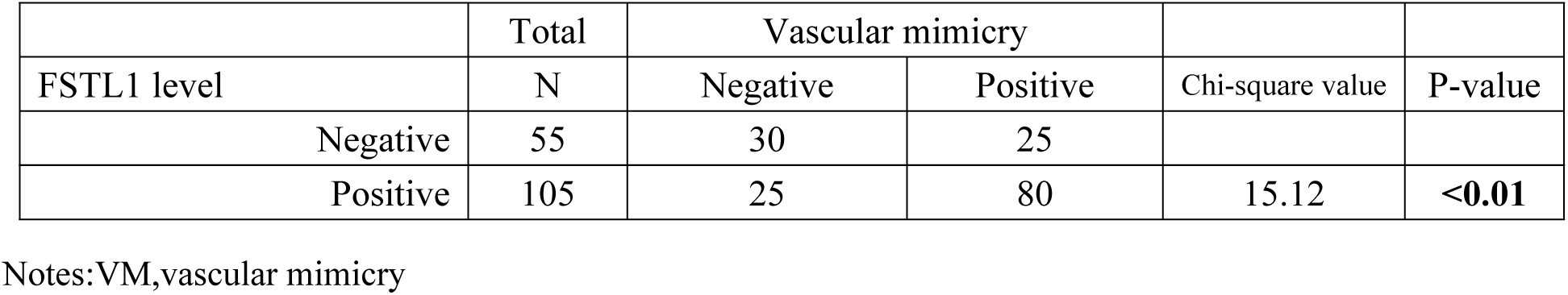
Association between FSTL1 levels and the presence of vascular mimicry.

### FSTL1 overexpression and knockdown influenced the malignant behavior of HCC cells

We measured the expression levels of FSTL1 in HepG2 and MHCC97-H HCC cells (Fig. 2A). The successful overexpression and silencing of FSTL1 using an FSTL1 plasmid in HepG2 cells and 2 targeted shRNAs in MHCC97H cells was confirmed via western blots (Fig. 2B). Similarly, we tested the proliferation, migration, and invasion of the two cell lines. We found that FSTL1 overexpression promoted increased proliferation, trans-Matrigel invasion, and colony formation of MHCC97H cells (Fig. 2C–E). Taken together, these findings indicated that abnormally high expression levels of FSTL1 in tumors may play a pro-tumorigenic role by promoting the proliferation, migration, and invasion of HCC cell lines. Additionally, qRT-PCR and western blot results revealed that FSTL1 promoted the expression of the EMT-promoting proteins Vim and Snail while downregulating the expression of PDCD4 and E-cadherin (Fig. 2F). FSTL1-shRNA inhibited the expression of Vimentin and Snail but promoted the expression of PDCD4 and E-cadherin. Notably, PDCD4 expression reversed the effects of FSTL1 overexpression. These results suggest that FSTL1 promotes increased blood supply to HCC tissues by regulating PDCD4.

**Fig. 2.**
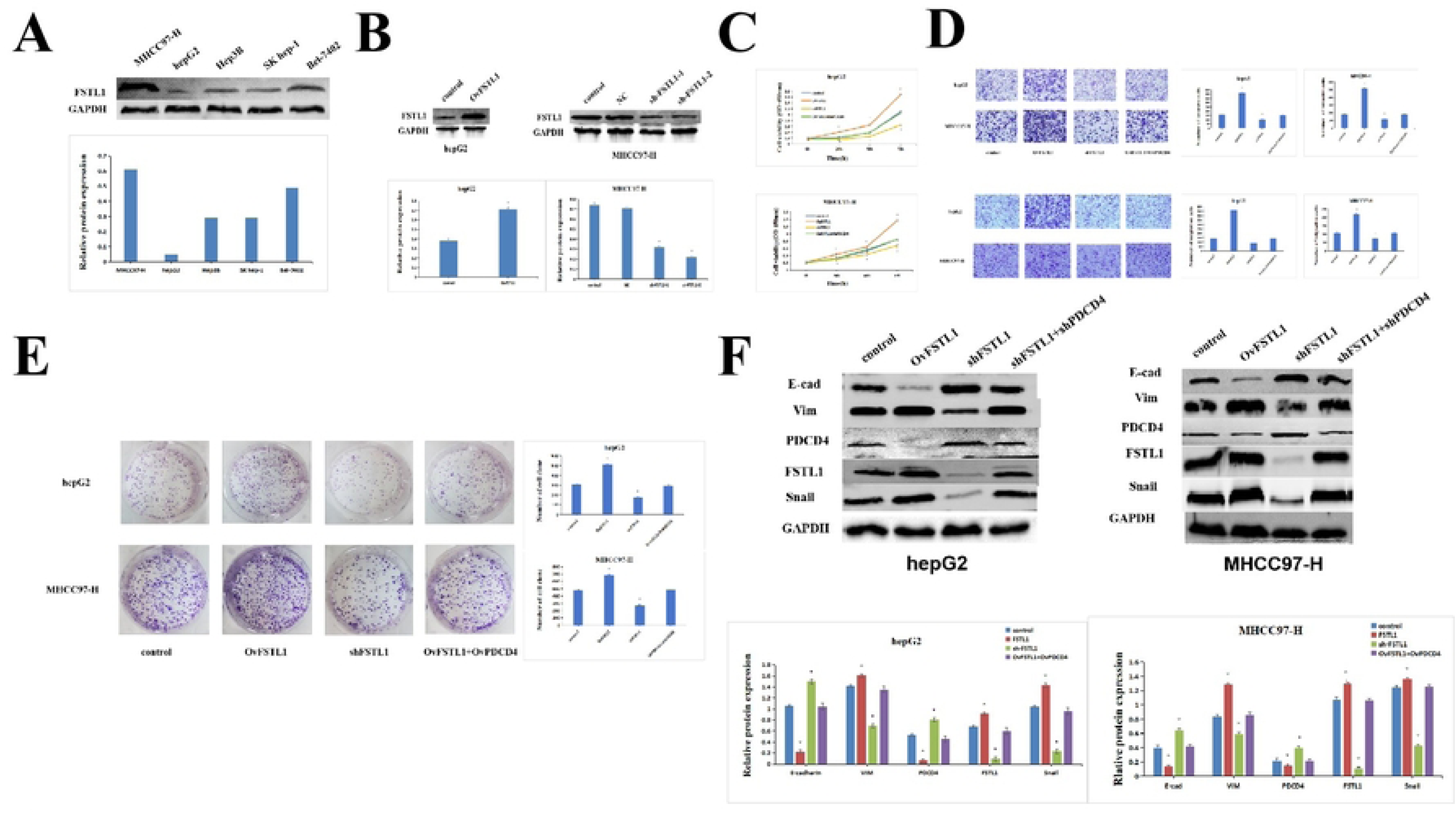
FSTL1 attenuates EMT of HCC cells by interacting with PDCD4. **A.**Western blot detected the expressions of FSTL1 in HCC cell lines. **B.** Western blot detected the expressions of FSTL1 hepG2 cells and MHCC97-H cells after the transfection **.C,E.** Proliferation ability of HCC cells was measured using CCK8 assays and colony formation assay after transfection. **D.** Transwell assay measured the migration and invasion ability of HCC cells. **F.** Western blot detects the expression of EMT-related proteins due to the interaction between FSTL1 and PDCD4. (E-cad, Vim and Snail-1).

### FSTL1 promotes VM in vitro

We next investigated the role of FSLT1 in vascular channel formation. As shown in Fig. 3A, FSLT1 expression led to a significant increase in tubular structure formation in HCC cell monolayers, whereas silencing of FSTL1 impaired their formation. The formation of these structures is thought to be associated with VEGF and HIF-1⍺ expression; therefore, we assessed the effect of FSTL1 and PDCD4 overexpression on HIF-1⍺ and VEGF levels. Western blotting and qRT-PCR revealed that FSTL1 promoted the upregulation of both VEGF and HIF-1⍺ in HCC cells, while PDCD4 inhibited this effect of FSTL1 (Fig. 3B). These results suggest a role for FSTL1 in tubular structure formation in vitro.

**Fig. 3.**
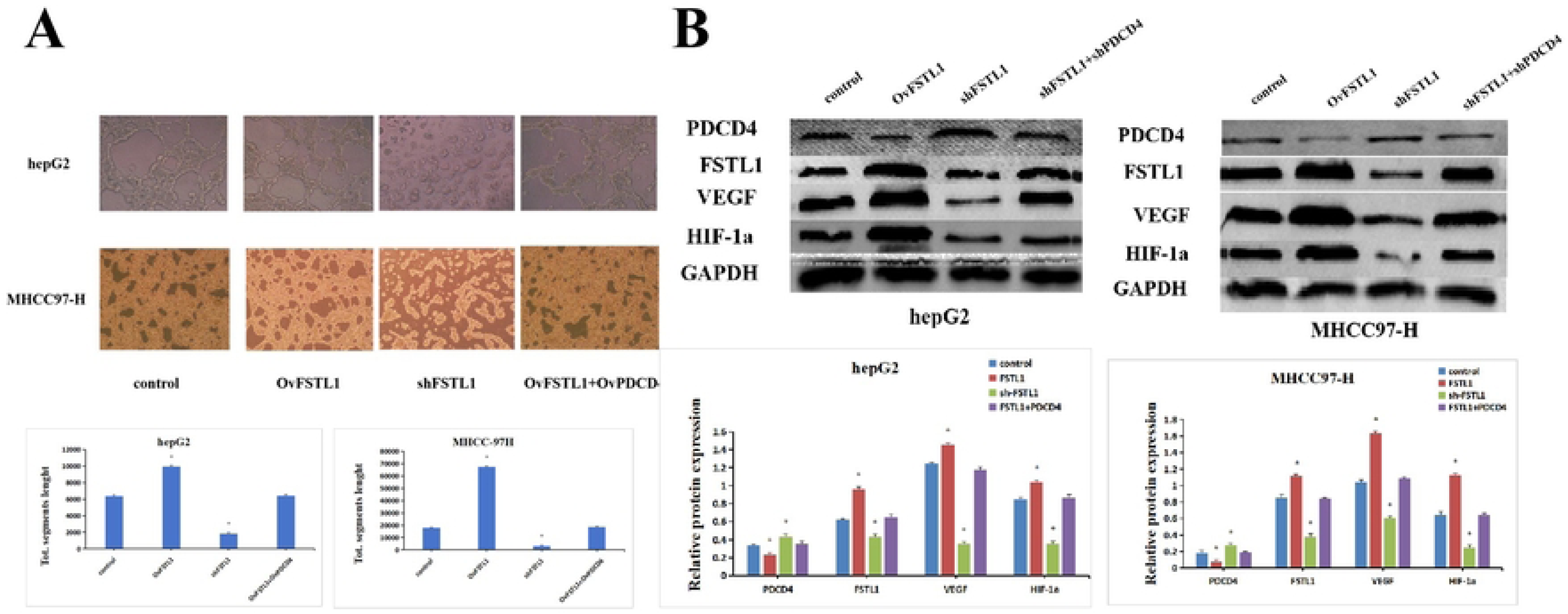
FSTL1 attenuates HCC cell angiogenesis by interacting with PDCD4.**A** Matrigel-based tube formation assay assessed the angiogenesis ability.**B** Western blot detects the expression of angiogenesis related proteins due to the interaction between FSTL1 and PDCD4. (VEGF and HIF-1a).

### FSTL1 activates the PI3K/Akt/mTOR pathway by inhibiting PDCD4 to promote VM in HCC cells

Our data showed that FSTL1 promotes the formation of VM in HCC cells by regulating PDCD4. Knockdown of FSTL1 effectively reduced the levels of phosphorylated PI3K, Akt, and mTOR in HepG2 and MHCC97H cells, while introducing PDCD4 restored their expression levels (Fig.4). These data suggest that FSTL1 promotes VM formation in part by activating PI3K/Akt/mTOR signaling in HCC cells

**Fig. 4.**
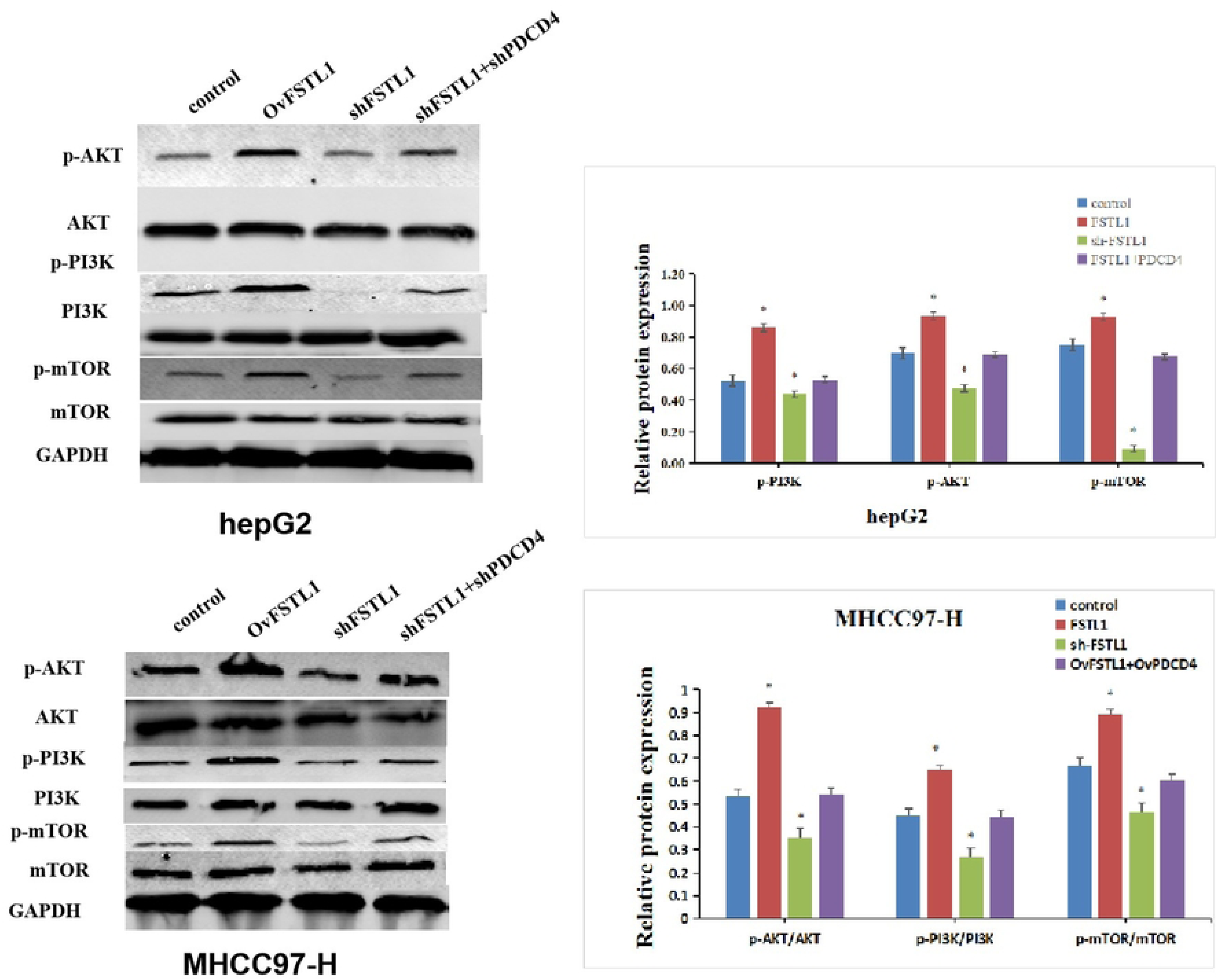
Effect of the interaction between FSTL1 and PDCD4 proteins on the expression of proteins related to the PI3K/Akt/mTOR pathway.

### FSTL1 activates the PI3K/Akt/mTOR pathway by inhibiting PDCD4 to promote VM formation in vivo

To further investigate the effects of FSTL1 on VM formation in vivo, we established MHCC97H tumor cell xenografts in nude mice; either control or FSTL1-silenced cells were injected into the flanks of each mouse (Fig. 5A), and the tumor volumes were measured every 3 days after day 13. As shown in Fig. 5B and 5C, tumors were significantly smaller in the FSTL1 knockdown group than in the control group (P<0.01). Furthermore, phospho-AKT, phospho-mTOR, and phospho-PI3K levels were lower in the FSTL1-knockdown group than in the control group; levels of vimentin and Snail were also significantly lower in the stable FSTL1-knockdown group, while levels of PDCD4 and E-cadherin were significantly higher (Fig. 5D).

**Fig. 5.**
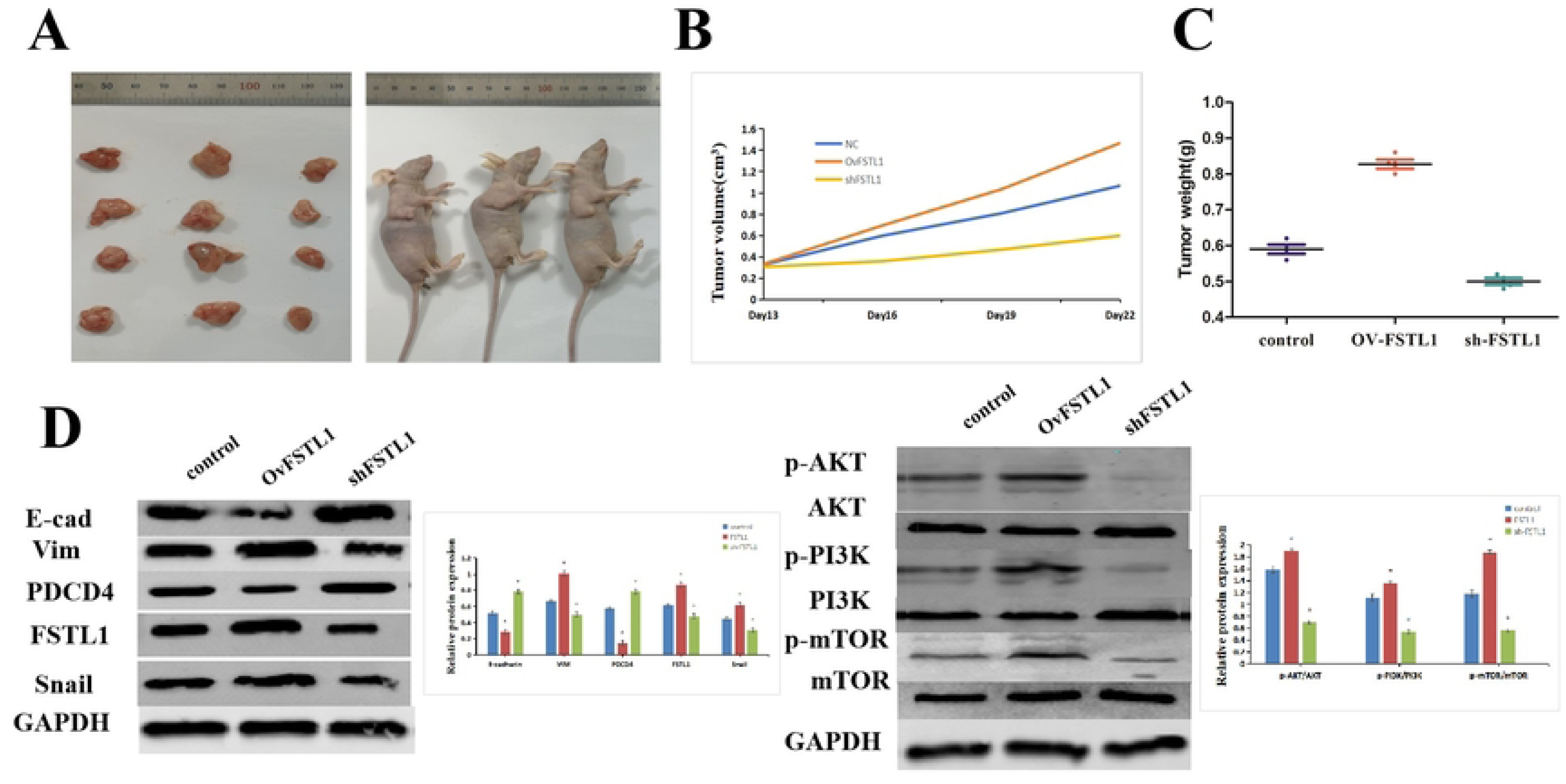
FSTL1 promotes the growth of transplanted tumors in nude mice by inhibiting PDCD4, regulates the expression of E-cad, vim, and snail, and activates the PI3K/Akt/mTOR pathway to promote VM formation in vivo.**A-C**.Tumor photos and tumor size throughout the experiment (22 days).**D**.Western blotting analyzed the impact of the interaction between FSTL1 and PDCD4 proteins on the expression of E-cad, vim, and snail proteins in tumor tissues, as well as the impact on PI3K/Akt/mTOR pathway-related proteins.E. Immunohistochemical analysis of the interaction between FSTL1 and PDCD4 proteins on the expression of angiogenesis-related proteins VEGF and CD31 in tumor tissues.

## Discussion

It is well known that the existence of blood vessels is a cornerstone of tumor cell growth and metastasis. VM, which is a precursor to blood vessel formation, can effectively boost such tumorigenic phenotypes. When existing endothelial cells cannot meet the requirements of tumor cells that are continuously growing, VM provides the necessary nutrients for these cells to maintain their unchecked proliferation (14). A hypoxic tumor microenvironment is an key potentiator of VM (15–17), as hypoxia activates pro-angiogenic factors such as VEGF and promotes the formation of blood vessels (18, 19). As such, VM can play a crucial role in the progression and metastasis of malignant tumors, and focusing on this phenomenon presents a novel research angle for understanding the formation and metastasis of tumors.

Previous studies have shown that FSTL1 plays an important role in ischemia-induced vascular remodeling in several tumor types including brain and central nervous system cancers (20) as well as colorectal (3), lung (21), and prostate cancers. In this study, we identified an important role for FSTL1 in the growth and metastasis of liver cancer cells in vitro and in vivo. Specifically, the interaction between FSTL1 and PDCD4 was found to regulate EMT in HCC cell lines. Additionally, FSTL1 was found to be involved in the activation of mTOR, which is a key kinase downstream of PI3K/AKT. The PI3K/AKT/mTOR pathway is an important signaling cascade that regulates tumor proliferation, growth, and blood vessel formation (22). More importantly, our analysis of patients’ clinical data revealed that high expression of FSTL1 in HCC tissues was closely associated with promoting invasion and other clinicopathological phenotypes.

The biological function of FSTL1 has been poorly understood to date. Although its involvement in some human malignancies has been demonstrated, the biological significance of FSTL1 as related to VM remains less clear. In this study, we first examined samples from 160 patients who had received definitive diagnoses of HCC and found that the expression of FSTL1 was closely related to tumor growth and metastasis. As the absorption of nutrients through blood is an essential condition for tumor growth and metastasis, we speculate that the high expression of FSTL1 may be related to VM, which is a precursor to neovascularization. Results of our CD31/PAS double staining and VM analyses were consistent with our hypothesis, as we found a large number of VM structures in HCC tissue sections that exhibited high expression levels of FSTL1. PDCD4 is a tumor suppressor that plays a crucial role in a variety of cellular functions such as controlling protein synthesis and the transcription of certain genes as well as inhibiting tumor invasion and metastasis; our co-immunoprecipitation experiments demonstrated that this protein interacts with FSTL1. To further investigate the effects of FSTL1 on the biological function of HCC via its modulation of PDCD4, we overexpressed or silenced FSTL1 expression in HCC cells and found that FSTL1 overexpression promotes tumor growth and metastasis while overexpressing PDCD4 reverses these FSTL1-induced phenotypes. EMT is an important component of tumor angiogenesis, and our western blots revealed that FSTL1 overexpression increases the levels of the EMT-modulating proteins Vim and Snail while decreased PDCD4 and E-cadherin levels (Fig. 2F). Conversely, FSTL1-shRNA inhibited the expression of Vimentin and Snail but promoted the expression of PDCD4 and E-cadherin. Given that PDCD4 overexpression inhibited the function of FSTL1, our results suggest that FSTL1 promotes vascular structure formation in HCC cells by regulating PDCD4 in vitro.

To further examine the role of FSLT1 in vascular structure formation, we conducted in vitro microvascularization experiments using genetically modified HCC cells. As shown in Fig. 3A, FSLT1 overexpression led to a significant increase in tubular structure formation; this in turn was impaired when introducing FSLT1 siRNA. Given that the formation of tumor blood vessels is associated with VEGF and HIF-1⍺, we assessed the effects of FSTL1 and PDCD4 expression on HIF-1⍺ and VEGF levels. We found that FSTL1 promoted the expression of VEGF and HIF-1⍺ proteins in HCC cells, while PDCD4 expression reversed this phenotype (Fig. 3B). These results suggested that FSTL1 plays a role in tubular structure formation in vitro.

Taken together, our data implicated FSTL1 in promoting VM in HCC through its interaction with PDCD4. The PI3K/AKT/mTOR pathway plays a critical role in cancer; mTOR is a key kinase downstream of PI3K/AKT that regulates the formation of tumor blood vessels. To further explore the mechanism of FSTL1 modulation of VM, we silenced FSTL1 and found that this effectively reduced the levels of phosphorylated PI3K, Akt, and mTOR in HepG2 and MHCC97H cells. Conversely, PDCD4 induction restored the expression levels of these proteins (Fig. 4). These data strongly suggested that FSTL1 promotes VM formation in part by activating PI3K/Akt/mTOR signaling in HCC cells.

We next investigated the effects of FSTL1 on VM formation in vivo using mouse xenograft models. As shown in Fig. 5B and 5C, tumors arising from FSTL1-silenced cells were significantly smaller than those arising from control cells (P<0.01). Additionally, CD31 and VEGFA expression levels were markedly lower in FSTL1-knockdown xenografts than in control tumors (P<0.001) (Fig. 5E). Importantly, phospho-AKT, phospho-mTOR, and phospho-PI3K expression levels were lower in FSTL1-silenced tissues than in control tissues. Vimentin and Snail were also significantly lower in the stable FSTL1-knockdown group, while PDCD4 and E-cadherin were significantly higher (Fig. 5D).

In conclusion, we found that the high expression of FSTL1 in HCC tissues is closely related to increased invasion and other clinicopathological phenotypes. We revealed an important role for FSTL1 in the growth and metastasis of liver cancer cells both in vitro and in vivo. Specifically, the interaction between FSTL1 and PDCD4 was found to regulate EMT in HCC cell lines; moreover, FSTL1 was found to be involved in the activation of AKT and mTOR, which are key kinases downstream of PI3K. The PI3K/AKT/mTOR pathway is known as a critical cancer signaling cascade that can regulate tumor proliferation, growth, and blood vessel formation. We conclude that FSTL1 promotes HCC VM by activating PI3K/AKT/mTOR signaling through its interaction with PDCD4 protein.

## conclusion

We conducted an analysis of clinical data obtained from patients diagnosed with hepatocellular carcinoma (HCC) and identified a significant association between heightened expression of FSTL1 in HCC tissues and the presence of invasive clinicopathological lesions. Our investigation revolved around comprehending the function of FSTL1 in the growth and spread of liver cancer cells, both in laboratory settings and in live organisms. We made an intriguing observation regarding the crucial role played by the interaction between FSTL1 and PDCD4 in the regulation of epithelial-mesenchymal transition (EMT) within HCC cell lines. Moreover, we noted the involvement of FSTL1 in the activation of the AKT/mTOR pathway, a critical downstream kinase pathway of PI3K/AKT. The PI3K/AKT/mTOR pathway is a well-established signaling pathway connected to cancer, responsible for controlling tumor proliferation, growth, and angiogenesis. To summarize, our discoveries strongly suggest that FSTL1 enhances hepatocellular carcinoma’s angiogenic mimicry by stimulating the PI3K/AKT/mTOR signaling pathway through its interaction with the PDCD4 protein.

## Abbreviations

Not applicable

## Ethics approval and consent to participate

Ethics Committee of China-Japan Friendship Hospital of Jilin University(2020-NSFC-074). Institutional Animal Care and Use Committee of Jilin University(KT202402255).

## Consent for publication

There no competing interests from involved authors all participants have given consent publish.

## Availability of data and materials

All data generated or analysed during this study are included in this article. We have not used other data that have already been published. All the data presented in this article are original results derived from this study.

## Authors’ contributions

Authors’ contributions: All authors contributed substantially to this study.

Li Hang and Tan Wenjia contributed equally to the article, and Li Hang is the co-corresponding author.

## Acknowledgements

Not applicable

